# A Novel Melatonergic Signature Predicts Reccurence Risk and Therapeutic Response in Breast Cancer Patients

**DOI:** 10.1101/472753

**Authors:** Hoa Quynh Tran, Phuc Loi Luu, Van Thai Than, Declan Clarke, Hanh Ngoc Lam, Dinh Truong Nguyen, Kim Van Thi Le, Trung Viet Nguyen, Minh Thong Le, Xuan The Hoang, Phan Q. Duy, Huyen Tran, Minh Nam Nguyen

## Abstract

ASMT is a key determinant of the levels of released melatonin. Though melatonin has been shown to exhibit anti-cancer activity and prevents endocrine resistance in breast cancer, the role of ASMT in breast cancer progression remains unclear. In this retrospective study, we analyzed gene expression profiles from thousands of patients and found that *ASMT* expression was significantly lower in breast cancer tumors relative to healthy tissue. Among cancer patients, those with greater expression had better relapse-free survival outcomes and longer metastasis-free survival times, and they also experienced longer periods before relapse or distance recurrence following tamoxifen treatment. Administration of melatonin, in combination with tamoxifen, further promoted cancer cell death by promoting apoptosis. Motivated by these results, we devised an ASMT gene signature that identifies low-risk cases with great accuracy. This signature was validated using both mRNA array and RNAseq datasets. Intriguingly, patients who are classified as high-risk benefit from adjuvant chemotherapy, and those with grade II tumors who are classified as low-risk exhibit improved overall survival and distance relapse-free outcomes following endocrine therapy. Our findings more clearly elucidate the roles of *ASMT,* provide strategies for improving the efficacy of tamoxifen treatment, and help to identify those patients who may maximally benefit from adjuvant or endocrine therapies.

## Introduction

Breast cancer constitutes the most common form of cancer in women (representing about 30% of all new cancer cases annually), and it is the second-leading cause of cancer deaths both worldwide and in the United States [1, 2]. About 75% of breast cancer patients are estrogen receptor (ER) positive, and the ER is a target for endocrine therapy such as tamoxifen administration [3–5]. Tamoxifen reduces recurrence rates (by ~39%) and mortality rates (by ~33%) in ER-positive breast cancer patients [4, 6–8]. However, among patients who initially respond to tamoxifen, 21-30% relapse within 4–14 years, even with 5 years of continued administration [4, 6, 7]. 30–50% of ER-positive breast cancer patients exhibit immediate resistance, whereas most patients develop resistance after initially responding to the drug [5]. Therefore, clinical resistance to tamoxifen presents a major challenge to successfully treating breast cancer.

Melatonin, which has been found to sensitize the response of breast cancer cells to tamoxifen both *in vitro* and in *vivo,* is primarily synthesized in the pineal gland [5]. It exerts both cytostatic and cytotoxic apoptotic effects in breast cancer by engaging the MT1 and MT2 receptors [5]. Tamoxfen efficacy can be improved by more than 100-fold in MCF-7 breast cancer cells with melatonin pretreatment [9]. It has also been shown to help in re-establishing the sensitivity of breast tumors to tamoxifen and tumor regression in a xenograft model [10]. Furthermore, supplementing tamoxifen administration with melatonin can slow the progression of metastatic breast cancer [11] and significantly increase response and survival rates in ER-negative metastatic breast cancer patients [12].

Acetylserotonin O-methyltransferase (ASMT), which plays roles in both healthy tissues and multiple cancer types, is a pivotal enzyme in melatonin synthesis. It has previously been combined with CYP1B1 as a two-gene prognostic index of human glioma [16]. However, the specific biological roles played by *ASMT* in cancer remain unknown. The relationship between *ASMT* expression and breast cancer risk (and survival outcomes) are poorly understood. As of yet, a clear understanding of the expression and prognostic potential of this gene in breast cancer and its clinical utility in endocrine therapy are lacking.

We investigate the significance of *ASMT* expression in human breast cancer and its relationship to survival outcomes. We find that low *ASMT* expression is a characteristic feature in breast tumors, and elevated levels of *ASMT* expression improve relapse-free survival (RFS) outcomes and metastasis-free survival times. Relative to other breast cancer patients, those with relatively higher expression levels of ASMT exhibit fewer relapses or longer distance recurrence outcomes following tamoxifen treatment. Administration of melatonin, in combination with tamoxifen, further promotes cancer cell death by upregulating apoptosis. Furthermore, we also devise a novel gene signature that can robustly predict recurrence risk. Patients predicted to be high-risk benefitted significantly from adjuvant chemotherapy. In addition, following endocrine therapy, low-risk patients benefited from better overall survival and distance relapse-free outcomes.

## Materials and methods

### Cell culture preparation

MCF7 cells were grown to 70% confluence in phenol-red-free RPMI 1640 supplemented with 10% fetal bovine serum, 100 units/ml penicillin and 100 μg/ml streptomycin sulfate at 37°C in a humidified 5% CO2 environment. Cells were pretreated with 1.0 mM melatonin for 2h then treated with 2.5 μM 4-OHT for 24h. Melatonin and 4-OHT were freshly prepared in a 1000X stock solution in DMSO and then diluted to the desired concentration directly in the culture medium. DMSO was added to control groups in each experiment. Live cells were observed under a Nikon Eclipse TE200 inverted microscope (Nikon, Tokyo, Japan).

### Cell apoptosis assays

The apoptotic cell distribution was evaluated using the Muse Annexin V & Dead Cell Kit from Merck Millipore (Danvers, MA, USA) per the manufacturer’s instructions. Following drug treatment, cells were collected and diluted with PBS containing 1% bovine serum albumin (BSA) to a concentration of ~5x10^5^ cells/ml. 100 μl of a single-cell suspension was mixed with 100 μl of Annexin V/dead reagent in a microtube and incubated in the dark for 20 minutes at room temperature (RT). Cells were then analyzed using the Muse cell analyzer from Merck Millipore (Danvers, MA, USA). The apoptotic ratio was determined by measuring the following four populations: (i) non-apoptotic cells: Annexin V(-) and 7-AAD(-); (ii) early apoptotic cells, Annexin V(+) and 7-AAD(-); (iii) late apoptotic cells, Annexin V(+) and 7-AAD(+); and (iv) died cells through non-apoptotic pathways: Annexin V(-) and 7-AAD(+).

### Colony-forming assays

Cells were seeded at 10^3^ cells/well in a 6-well plate and incubated for 24h. The cells were treated with DMSO, melatonin, 4-OHT or a combination of melatonin and 4-OHT for 6h. The media was then replaced by complete media. Cells were grown for 14 days to form colonies. Colonies were then fixed with acetic acid/methanol (1:7 v/v) at RT for 15 min. Crystal violet solution (0.5%) was added prior to 1h of incubation at RT, before washing with water.

### Western blot analyses

Cells were homogenized in RIPA buffer with protein inhibitors and centrifuged at 14,000 rpm for 10 minutes at 4°C. Protein concentrations were measured using the Bradford assay. Cell lysates (20 μg protein) were prepared in Laemmli buffer, heated for 5 minutes at 95°C, separated using sodium dodecyl sulfate-polyacrylamide gel electrophoresis (SDS-PAGE), and electroblotted onto nitrocellulose paper for 1h at 100 V. The presence of transferred proteins was assessed by Ponceau S red staining. The membranes were blocked with 3% BSA in TBST (20 mmol/L Tris-HCl, pH7.5, 50 mmol/L NaCl, 0.1% Tween 20). The membranes were then incubated for 1h with primary antibodies diluted to 1:1000 in TBST. After washing, the membranes were incubated for 1h with horseradish peroxidase-labeled secondary antibody diluted to 1:5000 in TBST, and the labeled proteins were detected using chemiluminescence reagents, following a protocol detailed by the manufacturer (Santa Cruz Biotechnology). Actin was immunoblotted to standardize the quantity of sample proteins for all western blotting analyses.

### Patients and gene expression profiles

Clinical and gene expression data were collected from GEO (http://www.ncbi.nlm.nih.gov/geo/), TCGA (http://www.cbioportal.org/index.do), and EMBL-EBI (https://www.ebi.ac.uk/arrayexpress/) under the following criteria: (1) sample sizes must be greater than 100, and (2) patients for whom data on disease progression are available were included (such as relapse time, survival status, etc.). Raw data were normalized as described previously [17]. 27 different datasets were used with a total of 7,328 samples (Supplementary Table 1). 635 samples were excluded, either because they were not tumors or because follow-up information was not available.

### The ASMT-associated gene signature

Datasets for building the *ASMT* gene signature were selected using the following criteria: (1) only patients with the status of RFS as patient endpoints were included, and (2) samples from these patients had to be assayed using Affymetrix HG-U133A chips. Three datasets (GSE1456, GSE2034, and GSE7390) with a total of 643 patients were used as a discovery set. Broadly, the gene signature was constructed in five steps (Fig. 4A). First, patients in the discovery dataset were separated into two groups (high- and low-expressing *ASMT* groups), based on the median *ASMT* expression. Second, a two-sample t-test identified genes that exhibit significant differential expression between the two groups (p<0.001). Third, a univariate Cox proportional hazard regression (p<0.001) was performed to identify the RFS-correlated genes. Fourth, common genes among different platforms were identified. Lastly, gene expression levels in the common gene set and RFS were used as features to build a survival risk classifier [18]. This classifier uses the principal component from the discovery dataset to produce a prognostic index for each patient. The prognostic index is computed as Σ*i*w*i*x*i*, where w*i* and x*i* designate the weight and log_2_-transformed gene expression for the *i*^th^ gene, respectively. The robustness of the classifier was evaluated using leave-one-out cross-validation (LOOCV). A patient was predicted to be high (low)-risk if their prognostic index was greater than (less than or equal to) the median prognostic index (0.782).

### Validation of the prognostic signature

Validation was performed on 23 independent mRNA array datasets (with 4,795 patients) and one RNAseq dataset from TCGA (with 1,096 patients). The median was subtracted from each gene individually. The Compound Covariate Predictor (CCP) was used as a class prediction algorithm to further refine this model and to sub-stratify outcome predictions [17, 19–21].

Kaplan-Meier survival analyses were performed after the samples were classified into two risk groups, and log-rank tests were used to evaluate risk. Uni- and multi-variate Cox proportional hazard regression analyses were conducted to evaluate independent prognostic factors. The gene signature, tumor grade, age, and molecular marker status were employed as covariates.

### Statistical methods of microarray data

We used the R language environment. A survival curve was analyzed and drawn using the Kaplan-Meier method, and p-values were evaluated using log-rank tests. The Cox proportional hazard regression model was used to carry out univariate and multivariate tests.

Cluster analyses were conducted with Cluster and Tree View [22]. P-values less than 0.05 were considered to be statistically significant.

## Results

### Upregulation of ASMTis associated with improved clinical outcomes

To test whether the *ASMT* expression is different between cancerous and normal breast tissue, we compared the mRNA in 270 normal and 395 breast tumor samples. On average, *ASMT* mRNA levels were significantly lower in tumor tissues (Fig. 1.A). To evalute the relationship between *ASMT* expression and clinical outcomes, we classified 2,529 patients into low and high expression groups (relative to the median *ASMT* expression). High expression of *ASMT* was positively correlated with better RFS outcomes (Fig. 1B). Notably, a Kaplan-Meier plot showed that breast cancer patients with high *ASMT* expression levels had longer metastasis-free survival (MFS) times (Fig. 1C). These results indicate that high expression of *ASMT* is not only positively related to reduced breast cancer risk, but is also associated with greater RFS and MFS times.

**Figure 1:**
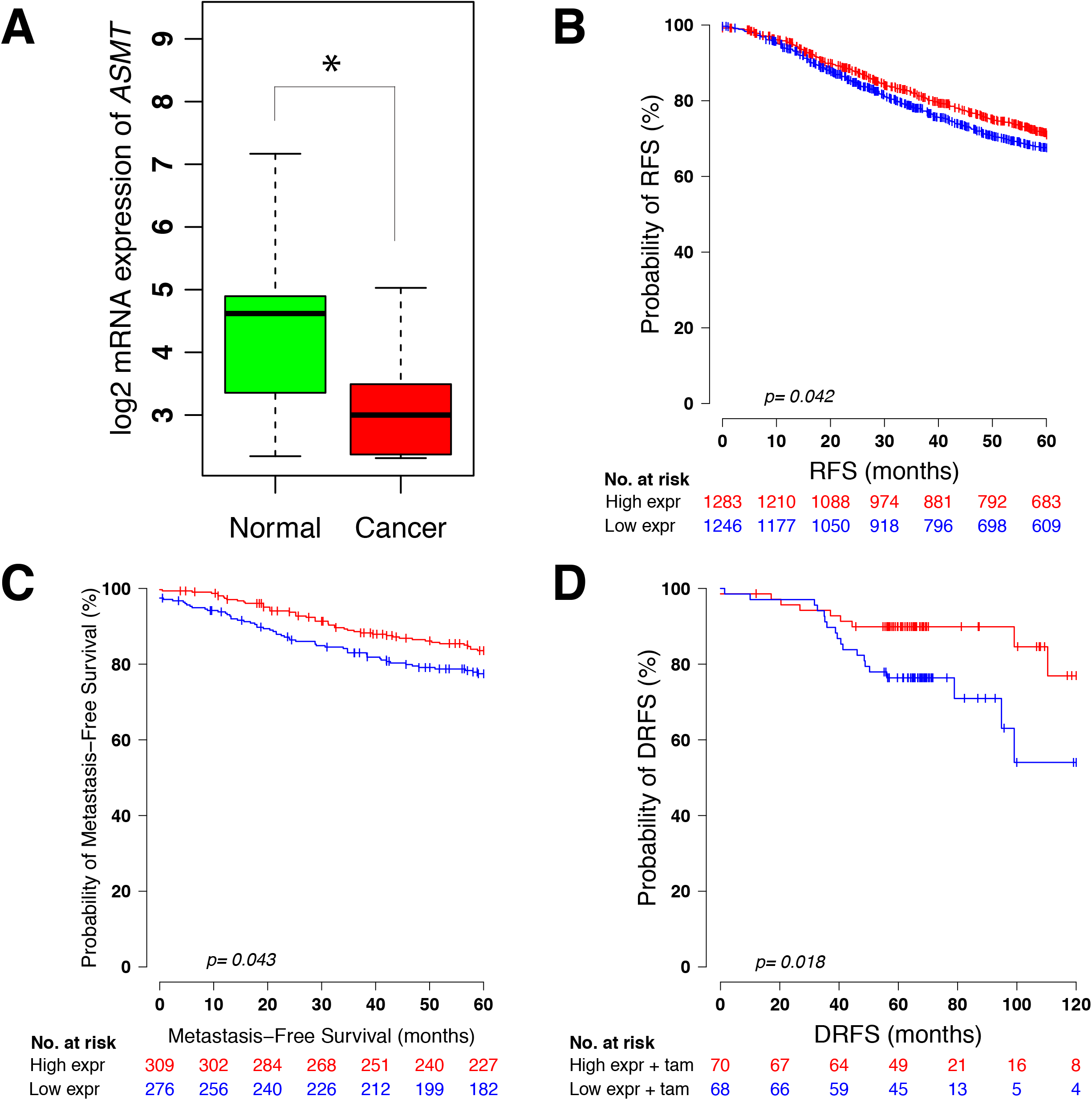
Upregulation of *ASMT* increases the survival rate and tamoxifen sensitivity in breast cancer patients. (A) Log_2_–transformed *ASMT* mRNA expression in normal and breast cancer tissues. Breast cancer patients were segregated into two groups according to their expression levels of *ASMT,* with median expression being used as the threshold to demarcate groups. Kaplan-Meier survival curves for that two groups were compared, with RFS (B) and metastasis-free survival. (D) A Kaplan-Meier plot for DRFS of that two groups of breast cancer patients who received tamoxifen treatment.

### Elevated expression of ASMT increases tamoxifen sensitivity

To investigate whether *ASMT* expression is related to tamoxifen treatment, we analyzed the GSE2990, GSE6532, and GSE9893 datasets which tamoxifen treatment availabe. Patients with relatively higher *ASMT* expression exhibited improved responses to tamoxifen treatment, and they exhibited longer survival times and distance recurrence after tamoxifen treatment (Fig. 1D and Fig.S1, respectively). *ASMT* expression thus promotes tamoxifen efficacy.

### Deep deletion of ASMT down-regulated protein expression of ER and HER2

To understand how ASMT expression promotes tamoxifen efficacy, we identified proteins which significantly changed by alteration of *ASMT.* We found that ER, HER2, AR, and PR were the top down-regulated proteins (Fig.S2). Protein expression of ER, ER-pS118, and HER2 were significantly decreased in patients had deep deletion of *ASMT.*

### Melatonin works synergistically with tamoxifen to promote cancer cell death

We evaluated whether dual tamoxifen-melatonin boosts sensitivity among patients for whom *ASMT* was lowly expressed. After MCF7 cells were treated with tamoxifen and melatonin, we observed that melatonin acts synergistically with tamoxifen to promote cancer cell death by elevating apoptositic pathways. The apoptosis markers (cleaved caspase 7 and cleaved PARP, Fig.3A) and the percentage of apoptotic cells were greatly elevated with combined melatonin-tamoxifen treatment, relative to melatonin or tamoxifen treatment individually (Fig. 3B). The combination treatment also strongly inhibits proliferation of MCF7 cells (as shown by the clonogenic assay in Fig. 3C).

**Figure 2:**
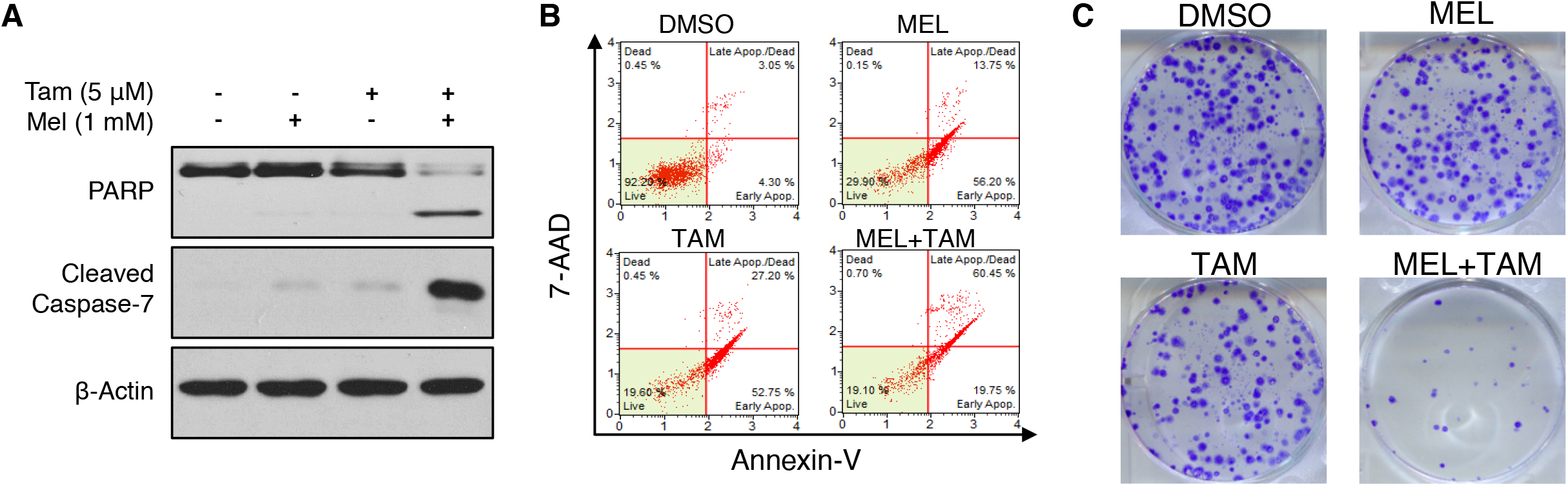
Synergiscal effect of melatonin and tamoxifen on cancer cell death. Representative images of western blot of apoptotic markers (A), cell apoptosis assay (B), and colony-forming assay (C) of MCF-7 cells after 24h of co-treatment with 1mM melatonin and 5μM tamoxifen.

**Figure 3:**
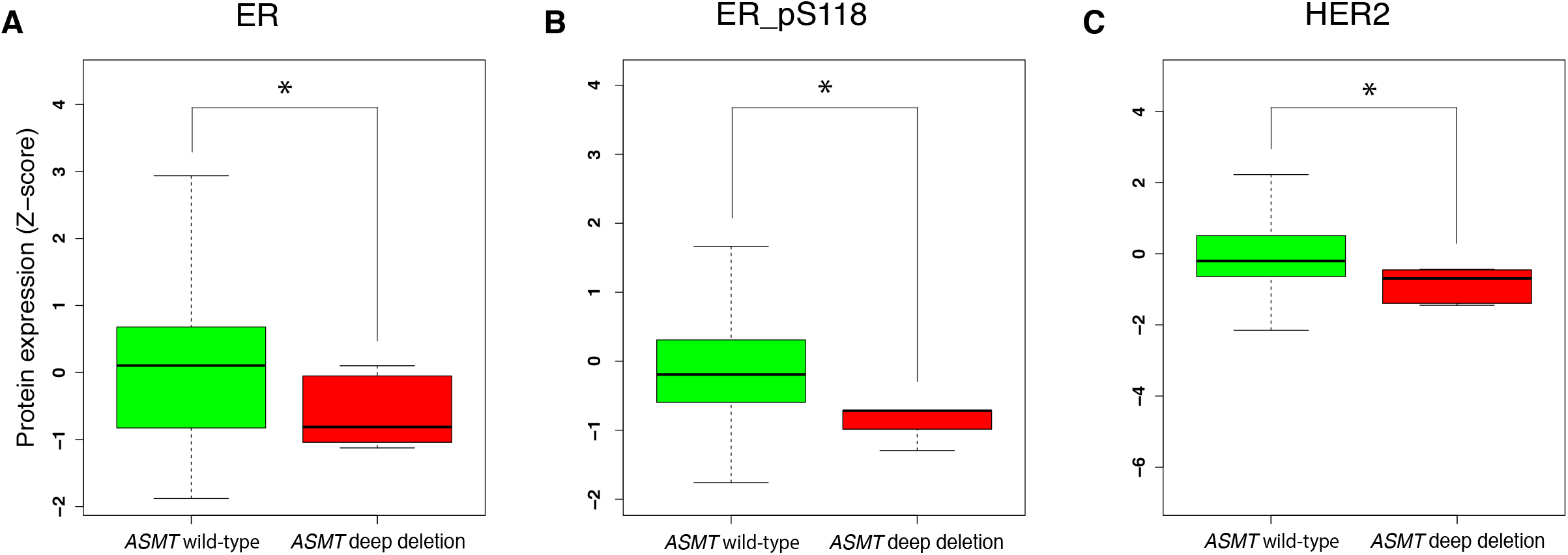
Deep deletion of *ASMT* downregulated ER and HER2. Boxplots showing the protein expression of ER (A), ER-pS118 (B), and HER2 (C) in breast patients with deep deletion of *ASMT* (n=6) relative to breast patients with *ASMT* wild-type (n_ = _838) normal controls (n_=_ 15). All P-values values were calculated using the two-sided Wilcoxon rank-sum test. (*) P_ <_0.05.

### Development of a ASMT signature for accurate risk prediction

Expression profiles were analyzed based on *ASMT* expression as it relates to empirical survival outcomes (Fig. 4 A). We identified 681 probes that were significantly differentially expressed between the high- and low-expression groups (Fig. S3). 37 genes were then identified using a survival univariate test (p<0.001). Finally, among these, 20 genes (which were shared between the different platforms) were selected as a gene signature and incorporated into a classifier based on a Cox proportional hazards model (Table S2). This 20-member set constitutes our melatonergic signature. We used the melatonergic signature to classify patients in the training set as high (n=322) and low (n=321) risk. Fig. 4B displays a heat map of the differentially expressed genes within the signature. There were significant differences between predicted groups in RFS (Fig. 4C), Overall Survival (OS; Fig. 4D), Distance Metastasis-Free Survival (DMFS; Fig. 4E), and Time to Distant Metastasis (TDM; Fig. 4F).

**Figure 4:**
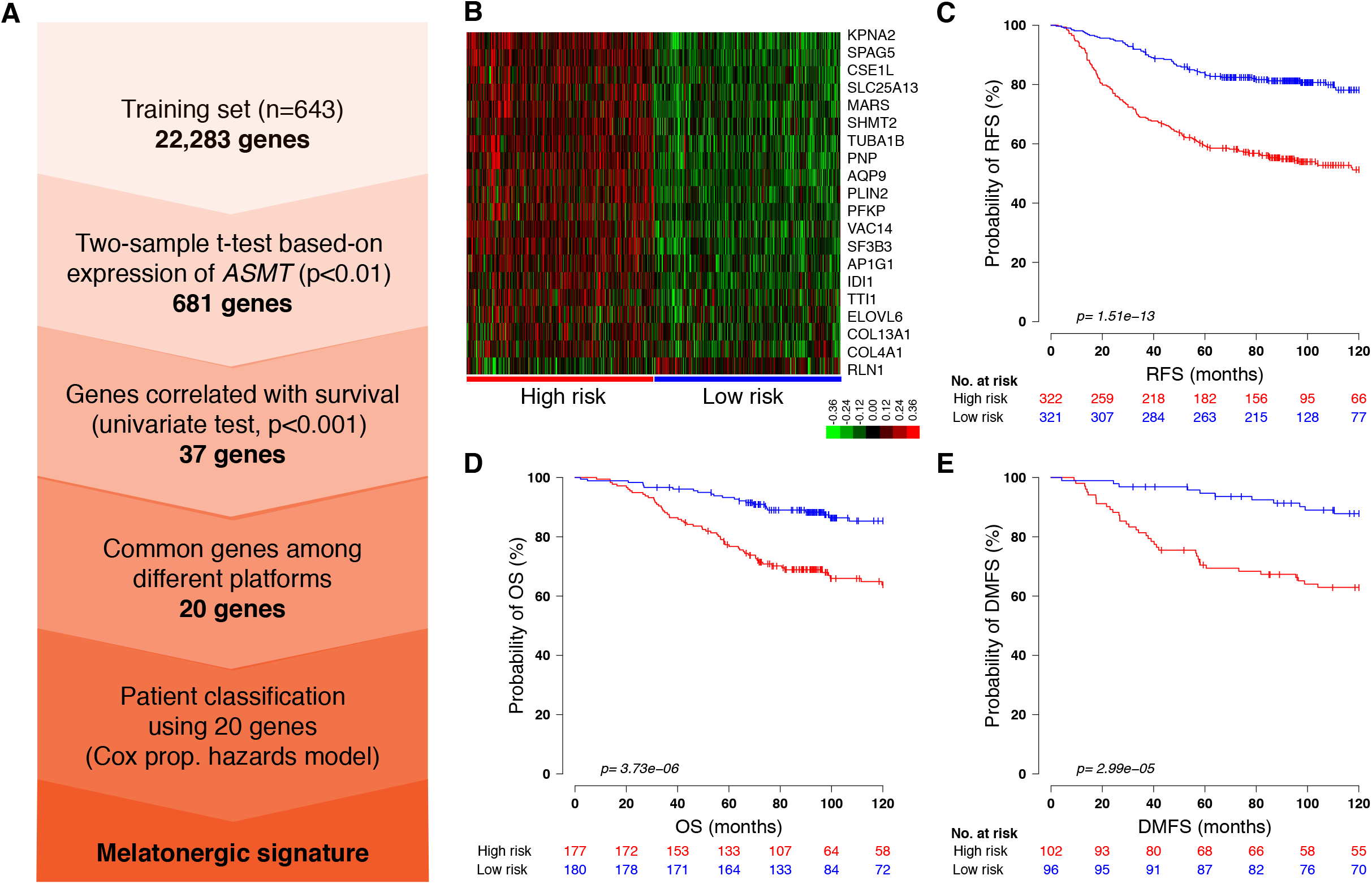
Discovering the genes associated with the melatonergic signature. (A) Schematic overview of the procedure used to construct the melatonergic signature based on gene expression profiles. (B) Hierarchical cluster analysis of the melatonergic signature in two risk groups of the discovery data set. Kaplan-Meier plots for RFS (C), OS (D), DMFS (E) and TDM (F) of the two risk groups in the discovery data set. P-values were computed using a t-test and a log-rank test.

### The Melatonergic signature accurately predicts and clinical features

To investigate whether the melatonergic signature is correlated with clinicopathological characteristics (such as age at diagnosis, tumor grade, histology, and follow-up time; Table 1). Tumor grade, ER status, histology, and follow-up times were significantly associated with melatonergic signature classification, whereas age showed no such association. Univariate and multivariate Cox regression analyses were performed on the discovery dataset to compare the prognostic value of the melatonergic signature with other prognostic covariates (Table 2). Interestingly, the melatonergic signature showed stronger prognostic predictive ability than these other clinical variables (Table 2). For both the univariate and the multivariate analyses, the melatonergic signature was significant in OS and RFS. These data suggest that the melatonergic signature is a significant predictor of DFS, OS, and DMFS, and it is generally independent of age, grade, and ER status.

**Table 1.** Clinicopathological features of breast cancer patients

| Variables |  | Total (%) | Low risk (%) | High risk (%) | *p* |
| --- | --- | --- | --- | --- | --- |
| Number of patients (%) |  | 643 | 321(49.9) | 322 (50.1) |  |
| Age | ≤46 | 100 (15.6) | 41(12.8) | 59(18.4) | 0.1794 |
|  | >46 | 98 (15.3) | 55(17.1) | 43(13.4) |  |
| Grade | 1 | 30(4.7) | 23(7.1) | 7 (2.2) | 3.27E-14 |
|  | 2 | 83(12.9) | 58(18.0) | 25(7.8) |  |
|  | 3 | 83(12.9) | 14(4.3) | 69(21.4) |  |
| ER | 0 | 141(21.9) | 24(7.5) | 117(36.3) | 2.2E-16 |
|  | 1 | 343(53.3) | 213(66.4) | 130(40.4) |  |
| Histology | Basal | 25(3.9) | 2(0.6) | 23(7.1) | 7.96E-15 |
|  | Her2 | 15(2.3) | 1(0.3) | 14(4.3) |  |
|  | LumA | 39(6.1) | 29(9.0) | 10(3.1) |  |
|  | LumB | 23(3.6) | 4(1.2) | 10(5.9) |  |
|  | Normal | 37(5.7) | 34(10.6) | 3(0.9) |  |
| OS | 0 | 275(42.8) | 157(48.9) | 118(36.6) | 4.56E-06 |
|  | 1 | 82(12.8) | 23(7.2) | 59(18.3) |  |
| DFS | 0 | 130(20.2) | 78(24.3) | 52(16.1) | 1.53E-04 |
|  | 1 | 29(4.5) | 6(1.9) | 23(7.1) |  |
| RFS | 0 | 431(67.0) | 258(80.4) | 173(53.7) | 5.93E-13 |
|  | 1 | 212(33.0) | 63(19.6) | 149(46.3) |  |

DFS, disease-free survival; DSS, disease specific survival; OS, overall survival; *p*-values were obtained using Fisher's exact test.

**Table 2.** Univariate and multivariate Cox proportional hazard regression analyses of clinical variables in the training dataset

| Variable | OS |  |  |  | DMFS |  |  |  | RFS |  |  |  |
| --- | --- | --- | --- | --- | --- | --- | --- | --- | --- | --- | --- | --- |
|  | Univariate |  | Multivariate |  | Univariate |  | Multivariate |  | Univariate |  | Multivariate |  |
|  | HR<br>(95% CI) | *p*-value | HR<br>(95% CI) | *p*-value | HR<br>(95% CI) | *p*-value | HR<br>(95% CI) | *p*-value | HR<br>(95% CI) | *p*-value | HR<br>(95% CI) | *p*-value |
| **Age >45 vs <=45** | 0.97<br>(0.53- 1.78) | 0.93 | 1.06<br>(0.57-1.97) | 0.85 | 1.06<br>(0.60-1.87) | 0.84 | 1.23<br>(0.68-2.21) | 0.49 | 0.90<br>( 0.58-1.41) | 0.64 | 0.96<br>(0.61-1.54) | 0.88 |
| **ER 1 vs 0** | 0.33<br>(0.18-0.59) | 2.98E-04 | 0.49<br>(0.24-1.00) | 0.05 | 0.40<br>(0.22-0.71) | 0.002 | 0.71<br>(0.36-1.39) | 0.31 | 0.63<br>( 0.40-1.00) | 0.05 | 0.88<br>(0.51-1.53) | 0.64 |
| **Grade 2 vs1** | 4.37<br>( 1.03 -18.60) | <0.05 | 2.89<br>(0.67-12.52) | 0.16 | 5.38<br>(1.28-22.69) | 0.01 | 3.37<br>(0.79-14.49) | 0.10 | 2.52<br>(1.06-5.95) | 0.04 | 2.48<br>(1.04-5.89) | 0.04 |
| **Risk low vs high** | 0.25<br>( 0.12-0.53) | 2.42E-04 | 0.27<br>(0.12-0.64) | 2.78E-03 | 0.26<br>(0.13-0.52) | 1.04E-04 | 0.28<br>(0.13-0.62) | 1.59E-03 | 0.36<br>( 0.22-0.50) | 4.51E-05 | 0.28<br>(0.16-0.50) | 1.27E-05 |

RFS, relapse-free survival; DMFS, distance metastasis-free survival; OS, overall survival; HR, hazard ratio; CI, Confidence Interval; *p*-values were obtained us a $\chi^2$ -test.

### The melatonergic signature is successfully validated using multiple independent datasets

To evaluate the robustness of the melatonergic signature, we performed validation using 23 independent breast cancer mRNA array datasets and one RNAseq dataset. The procedure used to validate the external datasets is shown in the flowchart in Fig. 5A. Using the CCP classifier, the sensitivity and specificity for correctly predicting low risk were 0.997 and 0.916, respectively. The melatonergic signature successfully classified patients in different platforms into the two-risk groups (Table S3). A Kaplan Meier plot showed a significant difference between the two risk groups in platform HG-U136A, HG-U133_Plus_2, MLRG Human 21K, Ilumina, Agilent, HG_U95Av2 and TCGA (Fig. 5B-E, F and Fig. S4-6). Overall, the melatonergic signature correctly classified a significant number of patients across independent platforms using both mRNA array and RNAseq data.

**Figure 5:**
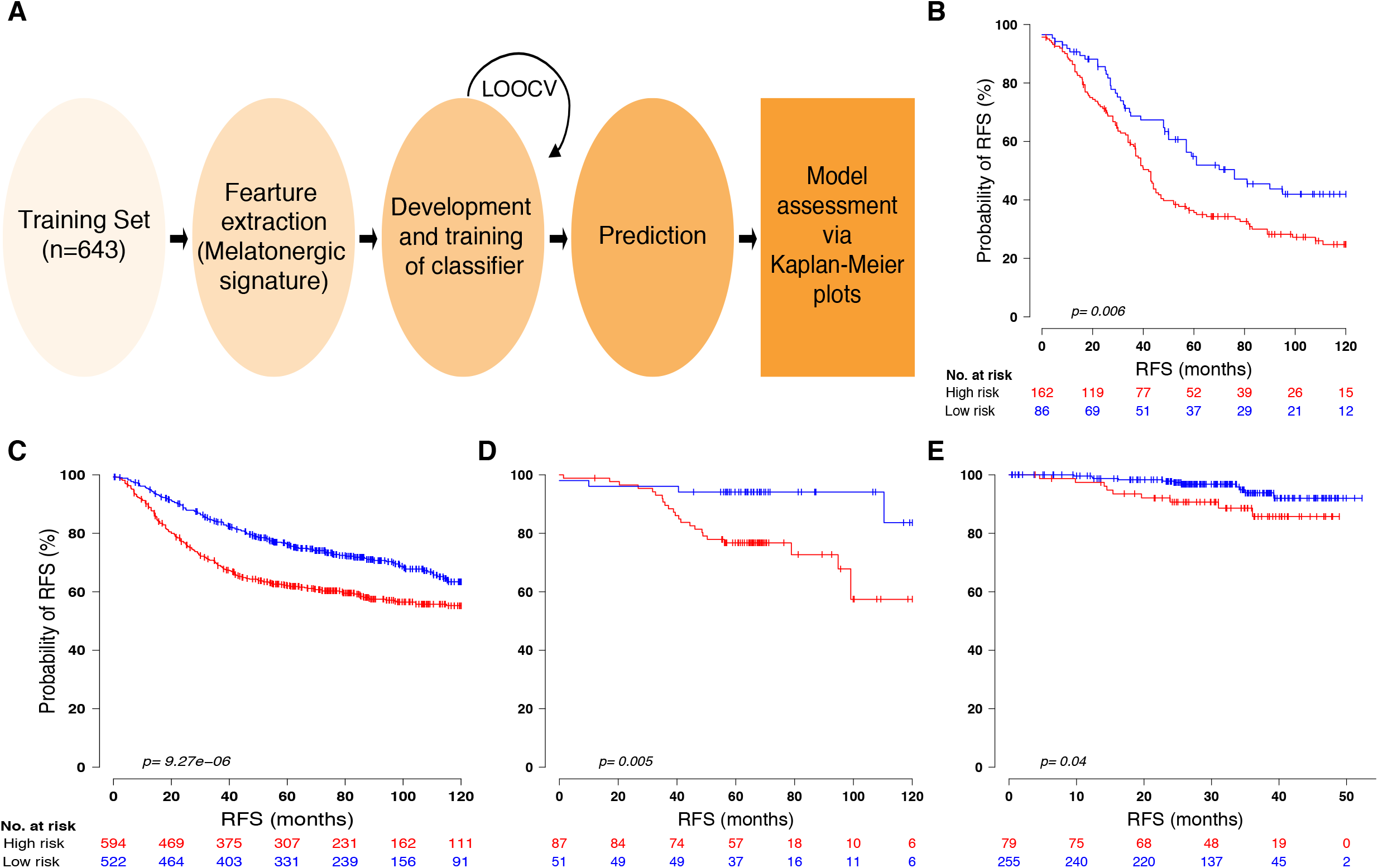
Prognostic significance of the melatonergic signature in independent validation cohorts. (A) Generating the risk prediction model and evaluating risk outcomes. Kaplan-Meier survival analyses for RFS of the validation datasets with the Human Genome U133A (B), Human Genome U133 Plus 2.0 (C), MLRG Human 21K V12.0 (D) and Illumina HumanRef-8 v3.0 expression beadchip (E) platforms. P-values were computed using a t-test and a log-rank test.

### The melatonergic signature correctly classifies patients with different tumor grades

A low-vs-high risk classification scheme can also be applied to patients with different tumor grades. The melatonergic signature accurately classified patients across all grades into low- and high-risk groups (Fig. 6 and table S4).

**Figure 6:**
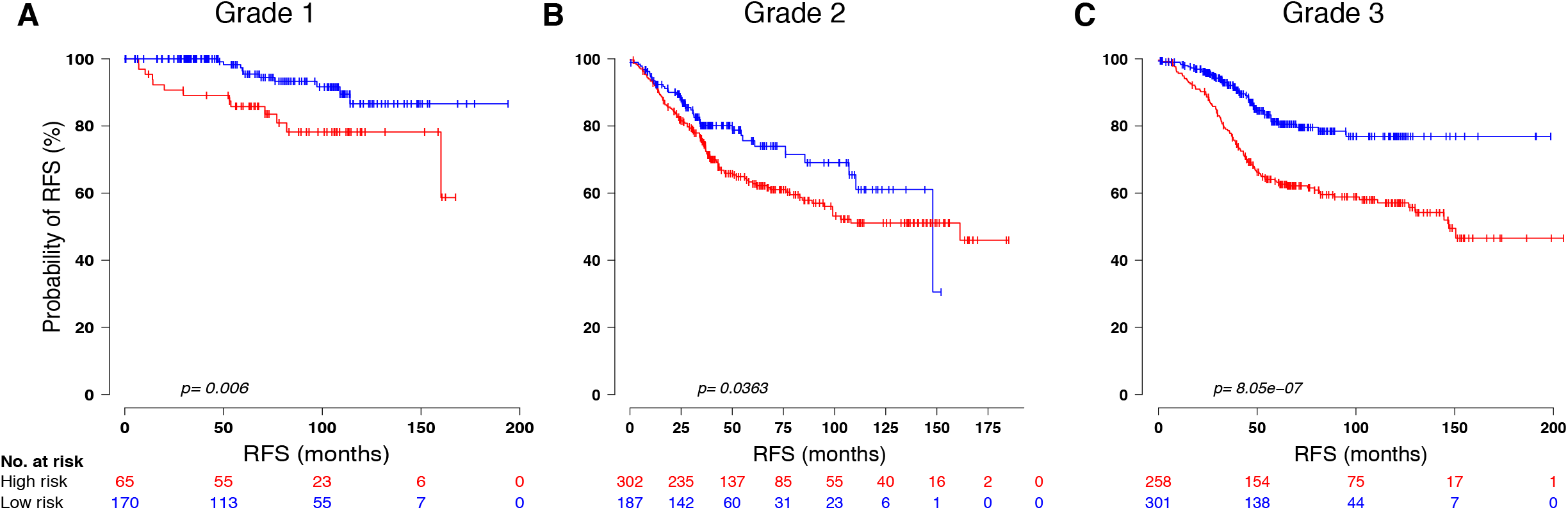
Association of the signature with different tumor grades. (A) Grade I, (B) Grade II, and (C) Grade III patients in validation datasets. The melatonergic signature was used to classify patients into high or low risk groups and evaluated by Kaplan-Meier survival analyses. P-values were computed using a t-test and a log-rank test.

### Leveraging the melatonergic signature to reliably stratify patients of different subtypes

A number of features associated with a particular cancer are often used to stratify breast cancer tumors into specific subtypes. However, considerable heterogeneity exists even within cancers that share the same features.

We thus investigated whether the melatonergic signature can further stratify breast cancer patients. We incorporated the melatonergic signature into ER, PR, HER, lymph node, and p53 statuses. The melatonergic signature was associated with ER-, HER-, PR+, PR-, Node+, Node-, and p53 wild-type (Fig. 7). However, we did not find significant differences between the two risk groups in ER-, HER+, and p53 mutation (Fig. 7). The melatonergic signature thus captures heterogeneity in patients of the same subtype, suggesting that it may help to overcome the limitations of the classification schemes using standard molecular markers.

**Figure 7:**
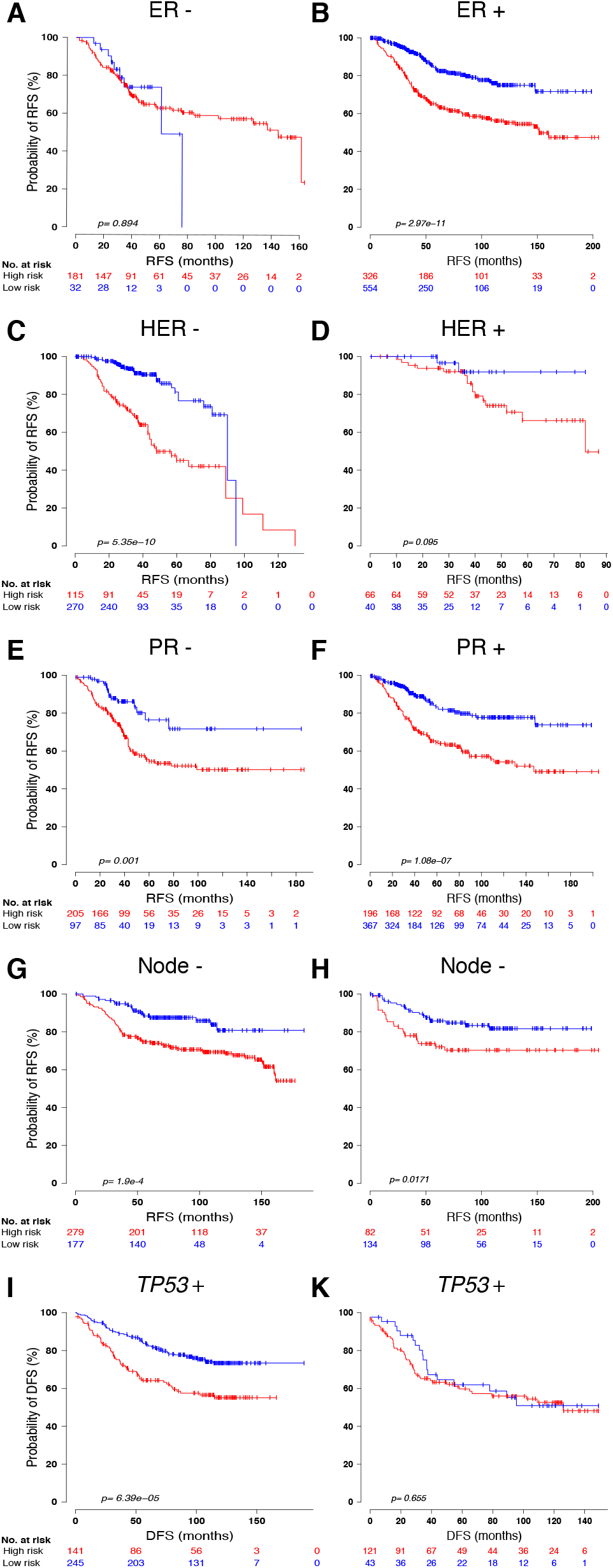
Prognostic significance of the melatonergic signature in molecular subtypes. Incorporation of the melatonergic signature into (A-B) ER, (C-D) HER, (E-F) PR, (G-H) Node, and p53 status of breast cancer patients. Patients were classified by the melatonergic signature into high and low risk groups and evaluated by Kaplan-Meier analyses. P-values were computed using a t-test and a log-rank test.

### The melatonergic signature can be used to better select patients for adjuvant chemotherapy and endocrine therapy

We evaluated whether the melatonergic signature may be used to identify patients that would optimally benefit from adjuvant chemotherapy and endocrine therapy. We selected patients who did not receive any treatment and patients treated with only adjuvant chemotherapy or endocrine therapy. To obviate confounding factors of potential synergistic effects, patients treated with a combination of both treatments (in addition to those treated with radiotherapy) were excluded. High-risk patients who received adjuvant chemotherapy exhibited more pronounced improvements relative to those who did not receive this treatment, though there was no significant differences among low-risk patients (Fig. 8A_B). Patients with low risk (in grade II) had better overall survival and distance RFS outcomes after endocrine therapy (Fig. 8C and 8E, respectively).

**Figure 8:**
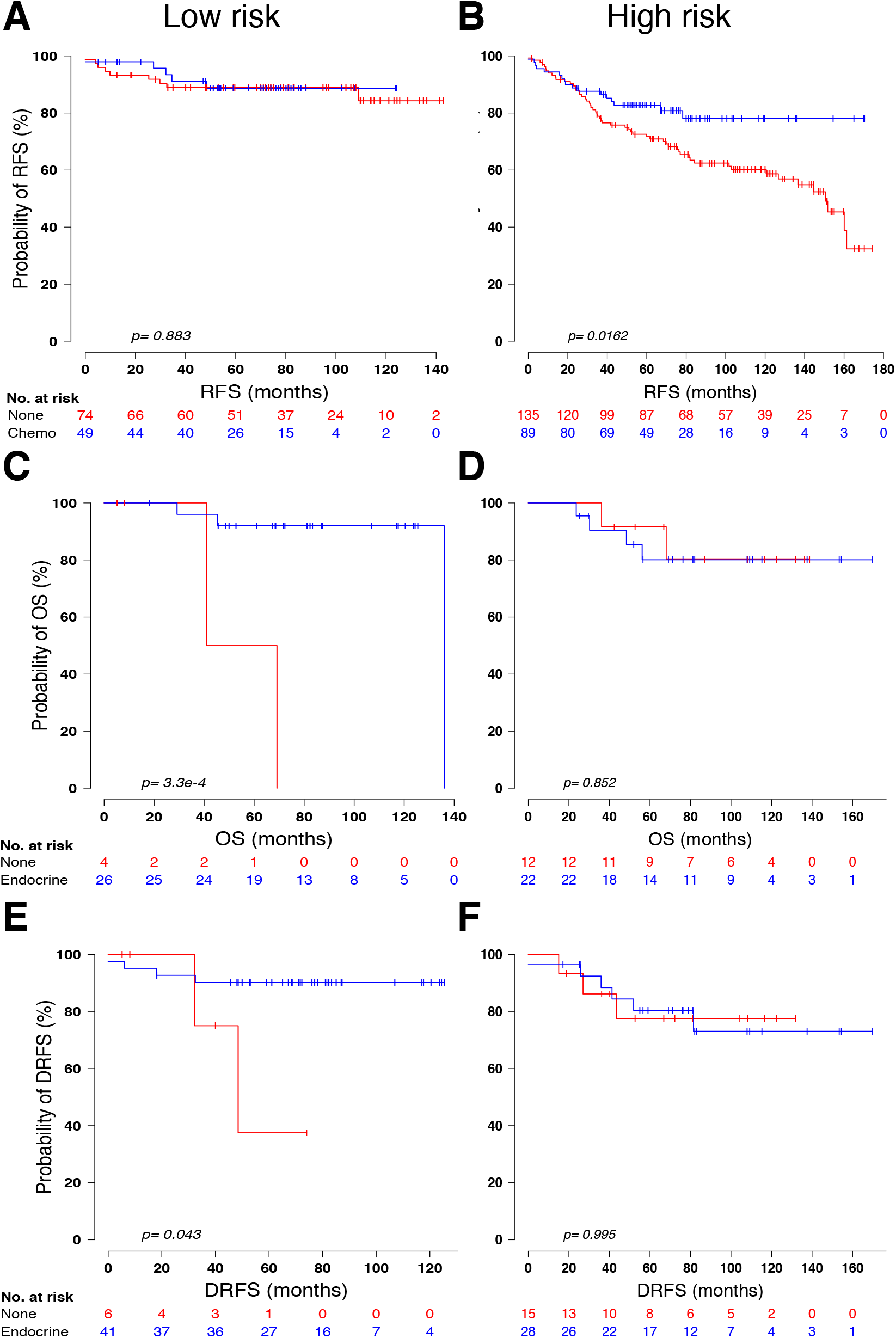
Kaplan-Meier survival analysis of the melatonergic signature and adjuvant chemotherapy and endocrine therapy. Patients were classified into different risk groups. Chemotherapy (A-B) or endocrine therapy (C-F), along with the conferred treatment, was evaluated by Kaplan-Meier analyses. P-values were computed using a t-test and a log-rank test.

## Discussion

The role of ASMT in cancer development remains unclear. Our work reveals that the expression of *ASMT* in breast cancer cells is significantly lower than that in healthy tissue. Breast cancer patients with relatively higher expression levels of *ASMT* exhibited improved RFS outcomes, longer MFS times, and better responses to tamoxifen treatment. Furthermore, melatonin was shown to promote cancer cell death by activating apoptotic pathways when combined with tamoxifen.

It has previously been reported that the function of the pineal gland is diminished and that the levels of melatonin are depressed in breast cancer patients [23]. It has also been reported that low levels of melatonin result in increased risk of developing breast cancer. Our study demonstrates that expression *ASMT* is diminished in breast cancer tissue relative to that in healthy breast tissue. It is likely that significantly impaired pineal functionality in breast cancer patients results in more moderate expression of *ASMT,* thereby resulting in diminished melatonin production and secretion. We emphasize that the expression of *ASMT* is significantly correlated with relapse-free and metastasis-free survival, suggesting the utility of quantifying *ASMT* expression for predicting clinical outcomes. Moreover, our results suggest that melatonin exerts a synergistic effect on inducing cell death via apoptotic pathways when combined with tamoxifen.

These results are thus consistent with previous experimental data demonstrating that melatonin inhibits the growth of MCF-7 cells [5], and that pretreatment of MCF-7 cells with melatonin augment the inhibitory actions of tamoxifen [9].

In a clinical trial, supplementing tamoxifen with melatonin has been found to prolong the pre-metastatic phase in patients with breast cancer [11]. In line with that finding, we found that patients with high *ASMT* expression levels tend to be more sensitive to treatment with tamoxifen and experience fewer relapses or longer phases before recurrence. We propose a model in which *ASMT* may sensitize cancer cells to tamoxifen, and that this gene may thus serve as a potential target for overcoming resistance in breast cancer patients.

In addition, for those patients in which the expression of *ASMT* was reduced, the proteins with the most significant reduction in expression were ER, PR, HER, and androgen receptor (AR). That may help to explain why tamoxifen exerts less of an effect in patients with reduced expression of *ASMT.* Melatonin stimulates ER expression, thereby significantly improving patient response to tamoxifen in ER-negative metastatic breast cancer [12]. Thus, we emphasize that, for cases in which breast cancer is ER-negative, *ASMT* may play a potential role in the improved responses to endocrine therapy. It may thus provide a much-needed tool for treating the difficult-to-treat cases in which breast cancer is ER-negative.

We also devised the melatonergic signature, which was validated in 5,891 patients. This signature indicated that high-risk patients may benefit from adjuvant chemotherapy, and low-risk patients (in grade II) have better overall survival and distance RFS outcomes following endocrine therapy.

The melatonergic signature consists of many genes which play important roles in migration/invasion, cancer survival, and proliferation of breast cancer cells (including *PFKP* [24, 25] and *COL4A1).* Several others (such as *AP1G1* and *SF3B3* [26, 27]) have been suggested as novel targets for overcoming drug resistance. *SHMT2, COL4A1, SF3B3, PLIN2, Elov16,* and *SPAG5* serve as independent prognostic factors in breast cancer patients, and high expression levels for these five genes were found to be significantly correlated with poor survival outcomes [27–33]. Many novel genes (such as *CSE1L, MARS, TTI1, VAC14,* and *TUBA1B)* were included in the melatonergic signature, suggesting that it contains new candidates which may play critical roles in breast cancer development. These may serve as novel diagnostic biomarkers and therapeutic targets for treatment.

Breast cancer is a heterogeneous disease with various subtypes. Though individuals with the same breast cancer subtype receive similar therapies [34], this heterogeneity results in only a subset of patients who substantively benefit from treatment [4–8]. The majority of breast cancers (60% to 75%) cannot clearly be classified into a particular subtype on the basis of these well-defined canonical features [35]. By integrating the melatonergic signature into the definitions of molecular subtypes, one can further sub-classify patients into two risk groups with greater resolution. The melatonergic signature may enable more informed therapeutic decisions and improved prognostics.

Histological grade and lymph node status are well-established prognostic factors [35]. [36, 37]. However, grade II invasive ductal carcinoma (without a clearly designated sub-type) presents considerable challenges in the clinical decision making process [38]. Although chemotherapy has been suggested to be especially advantageous for patients of grade III, there is no significant benefit in terms of DFS or OS with a single cycle of systemic treatment [37]. Furthermore, standard chemotherapy offers the same proportional reductions in recurrence and mortality among younger patients with node-positive or node-negative forms of the disease [7]. Thus, histological grade and lymph node status inadequately reflect heterogeneity. The melatonergic signature significantly and more finely stratifies patients in each histological grade and lymph node status group into prognostic subgroups.

The same therapy is generally applied to the majority of breast cancer patients [34], only a fraction of whom benefit from therapy [7, 39]. Our inability to precisely identify those patients for whom the disease will recur and those who respond to therapy stems from our limited understanding of disease heterogeneity. We demonstrate that patients identified to be high-risk significantly recur earlier than those identified to be low-risk, and that patients in the high-risk group may benefit from adjuvant therapy, whereas those in the low-risk group may be spared unnecessary and difficult treatment. Patients with grade II and low-risk have better overall survival and distance RFS outcomes after endocrine therapy. An estimated ~30% of all breast cancer patients are considered to be over-diagnosed and over-treated [40]. The melatonergic signature is a powerful tool that may help to obviate over-treatment and over-diagnosis.

The retrospective nature of our analyses represents one potential limitation. However, we note that this may help to mitigate the possibility of bias that may otherwise be introduced by experimental design. In addition, many novel genes in our signature have not previously been found to play a causal role in breast cancer progression.

This is the first study to document the critical role of *ASMT* in tumor progression in breast cancer, and it highlights the importance of *ASMT* both as a valuable patient-specific prognostic marker and as a potential target for personalized breast cancer therapy. We demonstrate that the melatonergic signature serves as a powerful predictor in gaining greater insights into the heterogeneous landscape of breast cancer, and that it can thus identify those patients who may maximally benefit from adjuvant and endocrine therapy. This signature has also identified number of novel breast cancer-associated genes. In addition to these promising new avenues for future study, this gene signature provides an accurate and easy-to-implement tool with general prognostic and clinical utility in the emerging age of precision medicine.

## Acknowledgments

We thank Emily Xiao (Yale University) for comments that greatly improved the manuscript. PQD was supported by NIH Medical Scientist Training Program Training Grant T32GM007205

---

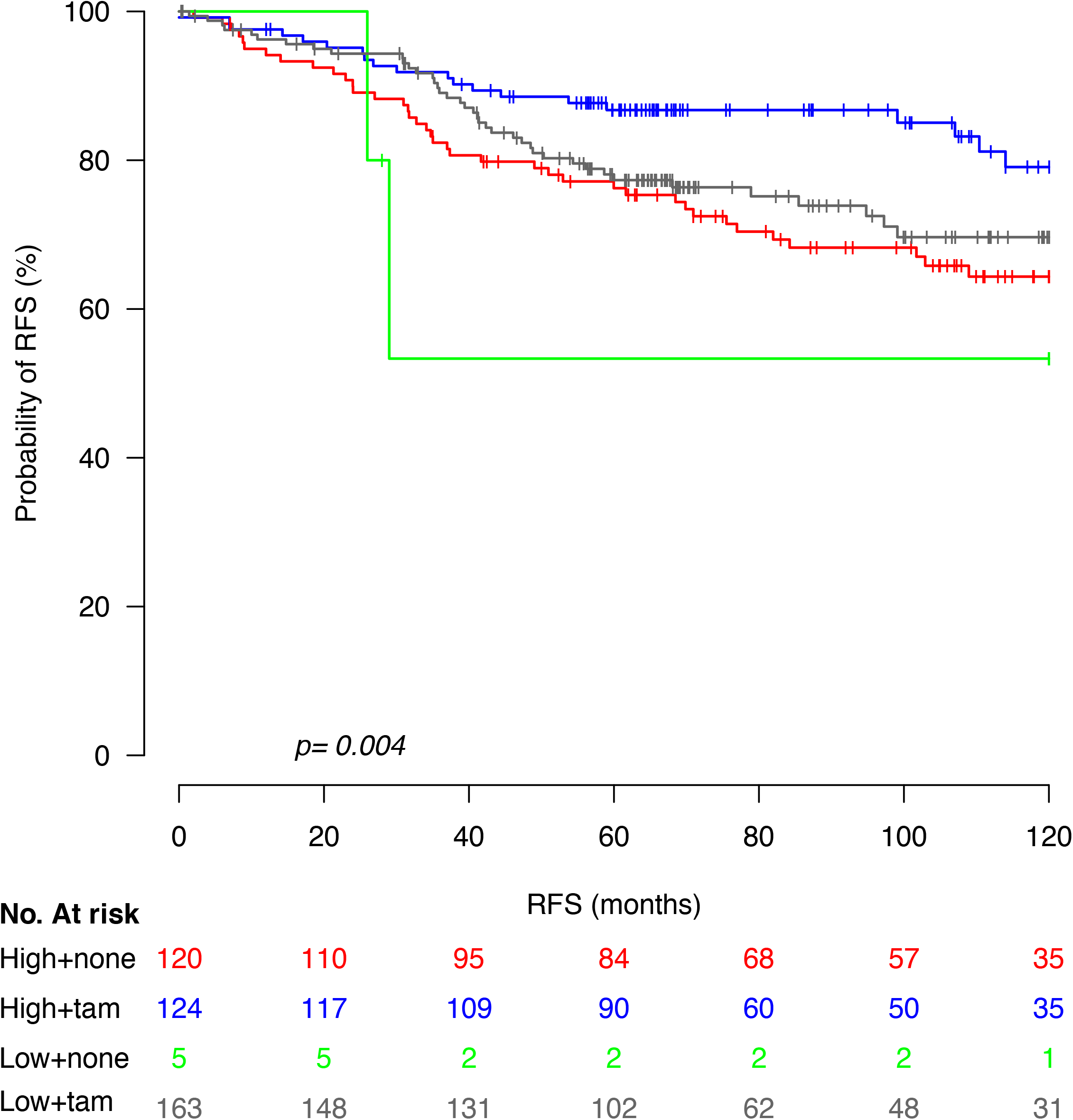

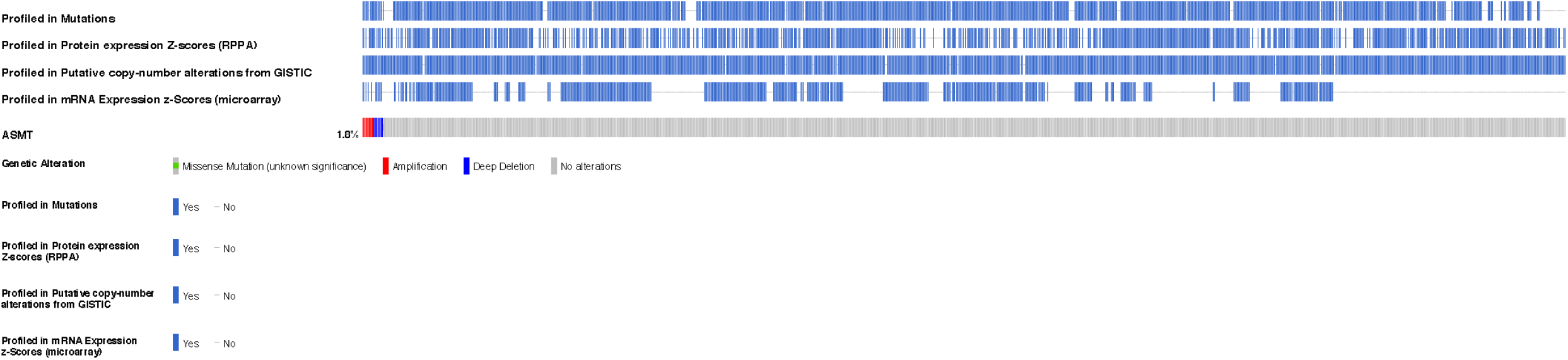

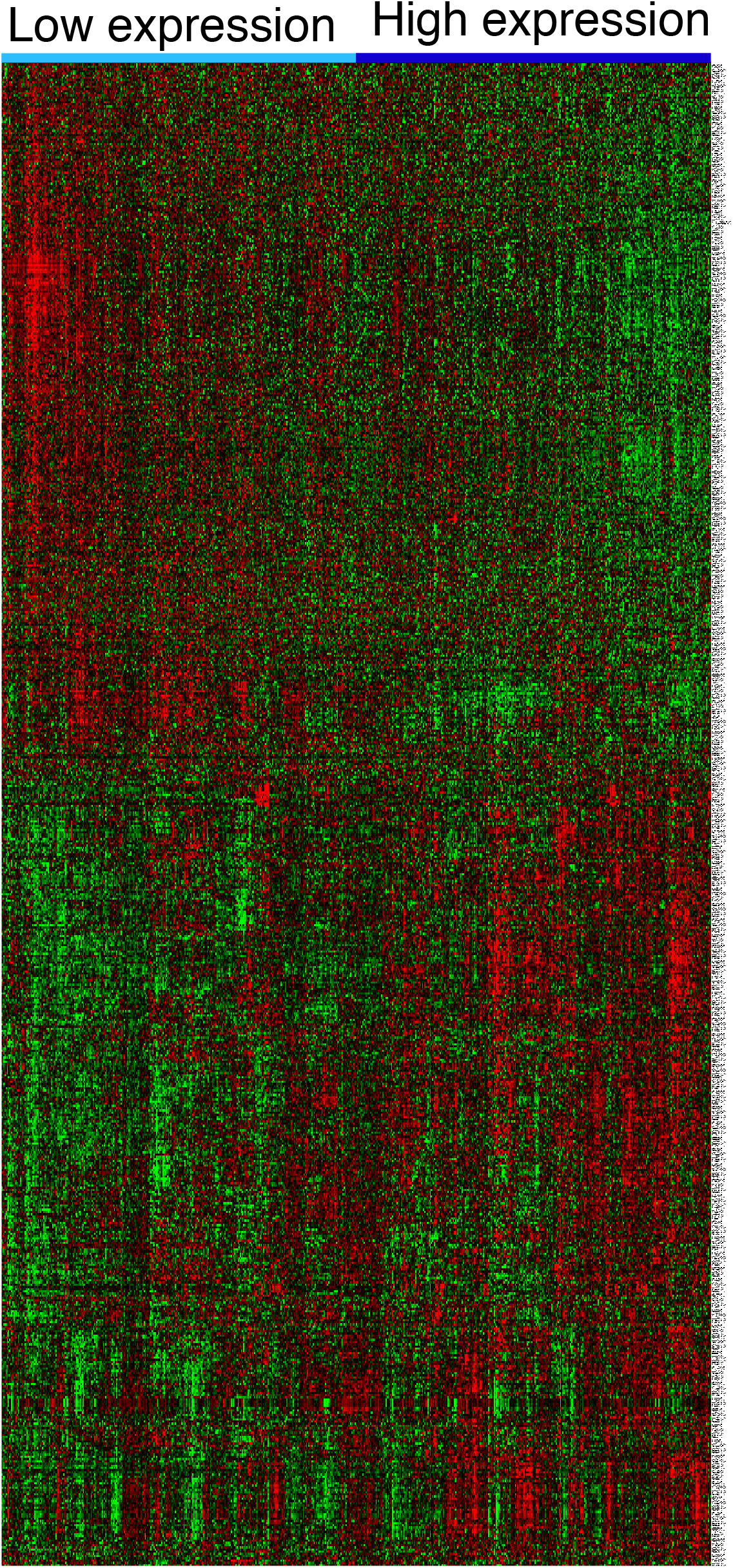

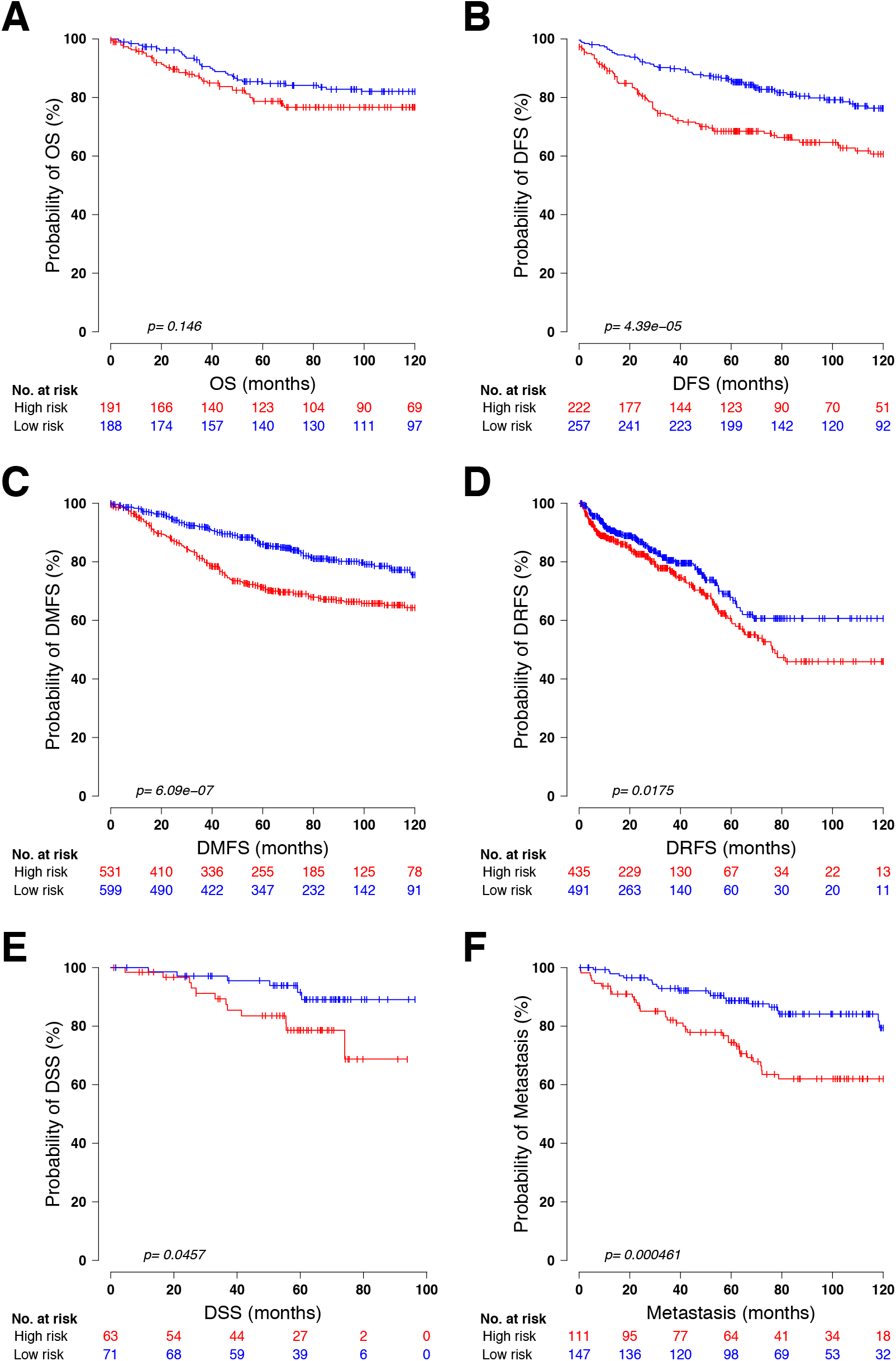

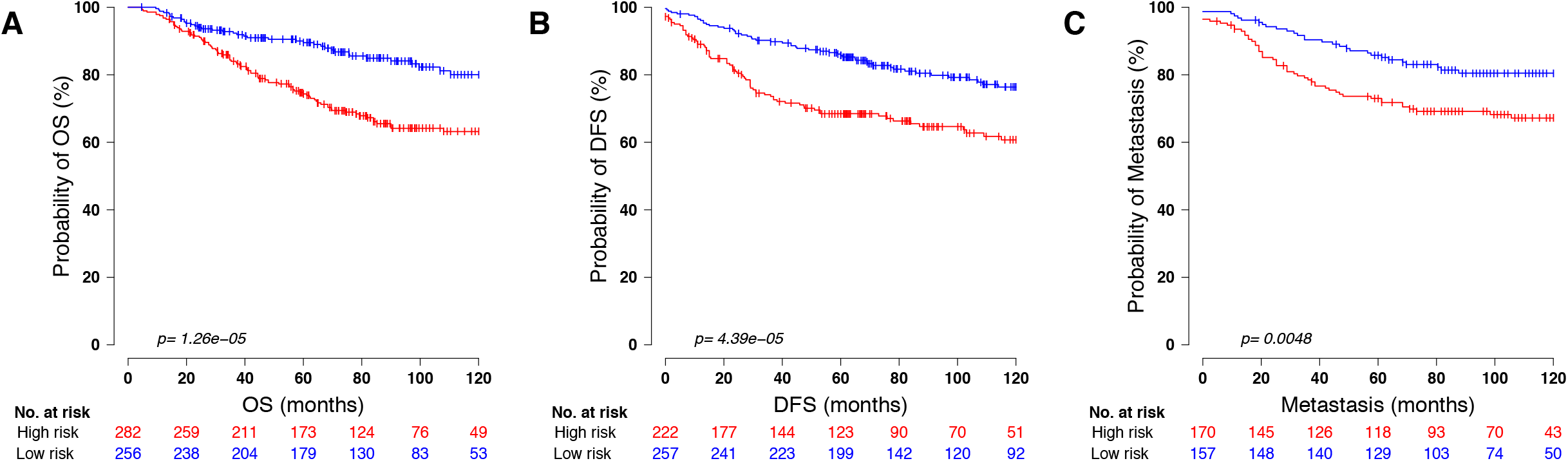

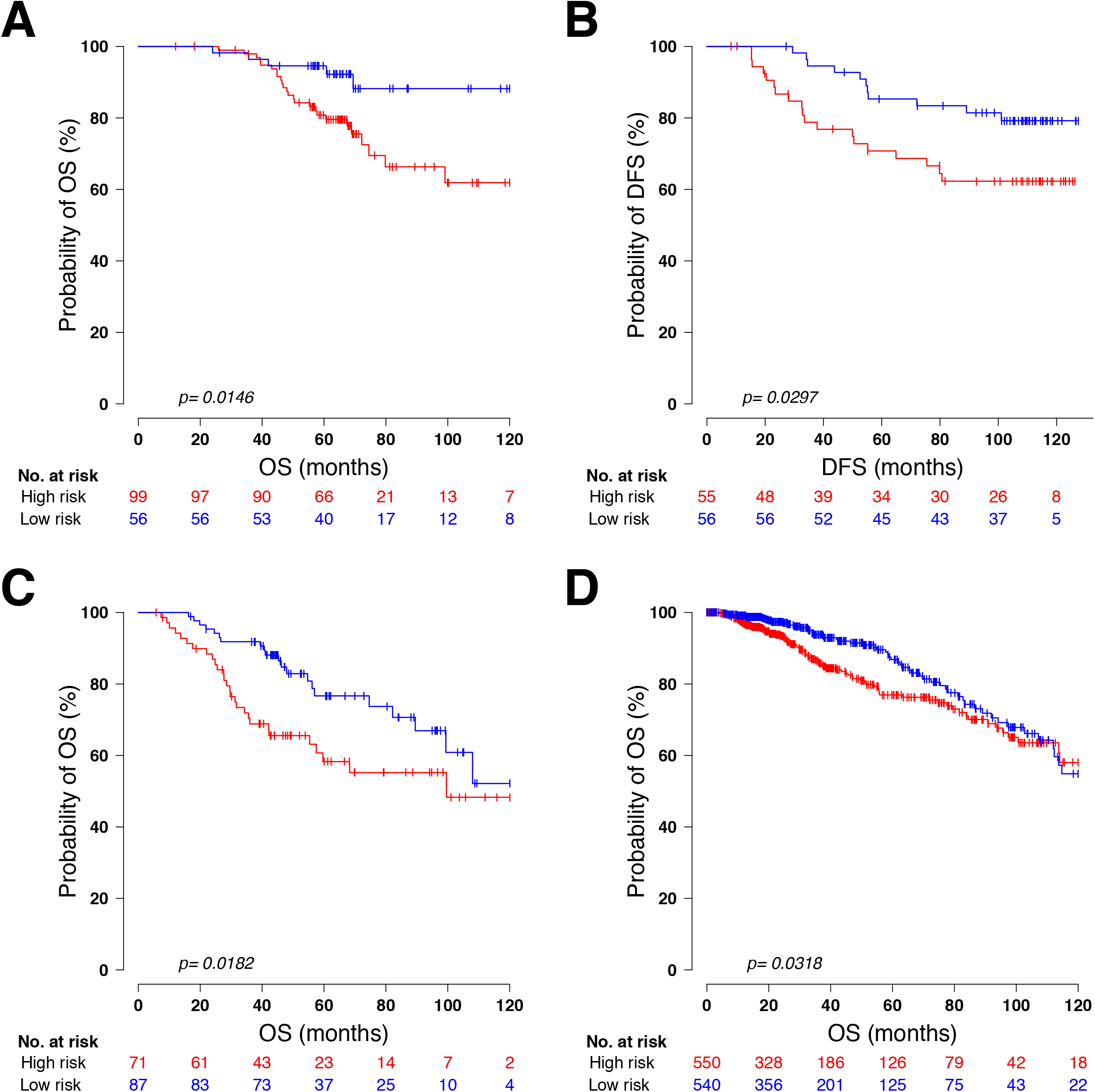

